# Exploration of the screening and regulatory mechanisms of biomarkers related to ac4C modification in laryngeal squamous cell carcinoma patients based on single-cell analysis and machine learning

**DOI:** 10.64898/2026.02.28.708684

**Authors:** Liqin Wang, Xiaoyang Gong, Donghui Chen, Xi chen, Han Zhou, Jiuhuang Lan, Renjing Ye, Zhuoding Luo, Yawen Shi

**Author notes:** Corresponding author: Yawen Shi, MS, Departments of Otorhinolaryngology, The First Affiliated Hospital, Nanjing Medical University, 300 Guangzhou Road, Nanjing 210029, China. These authors contributed equally (Liqin Wang, Xiaoyang Gong).

## Abstract

**Background:** N4-acetylcytidine (ac4C) modification plays a critical role in cancer development. Exploring ac4C modification in laryngeal squamous cell carcinoma (LSCC) may help elucidate its pathogenesis.

**Methods:** LSCC-related datasets were obtained from GEO. After preprocessing and annotating single-cell data, malignant cells were identified by CNV scoring and further divided into subpopulations. Malignant epithelial cells (MECs) were identified and subclustered based on ac4C-related gene activity. Prognostic genes were screened using Cox regression and machine-learning approaches, followed by validation in clinical samples using qPCR. The biological and immunological relevance of these genes was further explored through immune infiltration, immunotherapy response, and mutation analyses.

**Results:** The 14,465 identified MECs were classified into five subgroups (MEC1-5), among which MEC3 showed the strongest association with the ac4C gene set. Machine-learning analysis of MEC3-derived genes yielded seven prognostic markers, including BARX1, FHL2, NXPH4, PKMYT1, TNFAIP8L1, CRLF1, and CENPP. qPCR confirmed their differential expression between tumor and adjacent normal tissues. These genes were significantly associated with alterations in the tumor immune microenvironment, with high-risk patients showing increased immune infiltration and immune activity.

**Conclusion:** Seven ac4C-related prognostic genes were identified that may contribute to LSCC progression by modulating the tumor immune microenvironment, providing potential therapeutic insights.

## 1. Introduction

Laryngeal squamous cell carcinoma (LSCC) is the most common histological subtype of laryngeal cancer, accounting for approximately 85% to 95% of all cases[1, 2]. Its clinical management remains challenging due to the frequently nonspecific early symptomatology, resulting in over 40% of patients presenting with advanced-stage disease at initial diagnosis[3, 4]. The pathogenesis of LSCC involves a complex interplay of environmental exposures, viral oncogenesis, and genetic susceptibility factors[5]. While contemporary treatment modalities-including refined surgical techniques, precision radiotherapy, and evolving chemotherapeutic protocols-have demonstrated incremental improvements, the persistent challenges of tumor invasion, metastatic progression, and acquired chemoresistance continue to drive elevated locoregional recurrence rates and diminished long-term survival[6, 7]. Consequently, the identification and validation of molecular biomarkers with prognostic significance emerge as crucial prerequisites for developing more effective therapeutic strategies and improving clinical outcomes.

RNA modification is a process of chemically modifying RNA molecules, involved in various biological processes[8]. RNA modifications such as N6-methyladenosine (m6A), pseudouridine (Ψ), 5-methylcytosine (m5C), and N4-acetylcytosine (ac4C) are among the most prevalent types[9]. Among these, ac4C modification, as a conserved RNA modification, represents the only acetylation event in eukaryotic RNA[10]. ac4C modification is mediated by N-acetyltransferase 10 and serves essential functions in stabilizing mRNA and enhancing its translation[11]. In recent years, the regulatory functions of ac4C modification in the field of oncology have garnered widespread attention. Numerous studies have elucidated the molecular mechanisms linking ac4C modification to cancer progression, including key aspects such as cell proliferation, metastasis, metabolism, and chemotherapy resistance[12–14]. The unique expression of ac4C in diseases and its pivotal impact on cancer development establish it as a viable target for novel diagnostic and therapeutic strategies[15, 16].

This study employs an integrated multi-omics approach, combining bulk and single-cell transcriptomic profiling, to systematically delineate the mechanistic contributions of ac4C modification to LSCC pathobiology. Our analytical paradigm encompasses: (1) single-cell resolution identification of ac4C-enriched cellular subpopulations within the LSCC tumor microenvironment, (2) inference of intercellular communication networks and transcription factor regulatory circuits underlying ac4C-mediated functional effects, (3) machine learning-driven feature selection combined with survival analysis to establish robust prognostic signatures, subsequently validated through experimental confirmation, (4) construction and rigorous validation of a clinical prognostic model based on ac4C-related prognostic genes, and (5) comprehensive interrogation of associated mechanisms through immune contexture characterization and therapeutic vulnerability assessment. Collectively, these investigations provide novel insights into the epigenetic regulation of LSCC and establish a conceptual framework for developing ac4C-targeted therapeutic modalities. The study flowchart is presented in Fig. 1.

**Figure 1.**
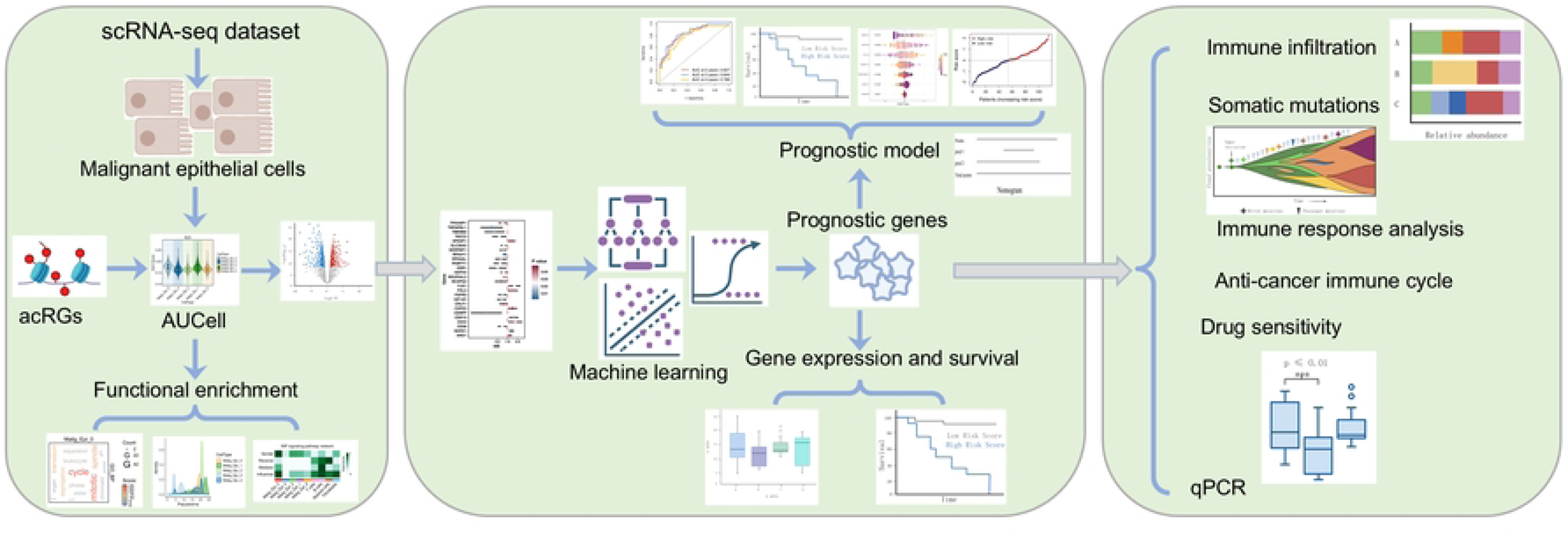
Study workflow for identifying ac4C-related prognostic biomarkers in LSCC.

## 2. Materials and methods

### 2.1 Data source

The LSCC-related datasets were obtained from the Gene Expression Omnibus (GEO) database (https://www.ncbi.nlm.nih.gov/geo/) and The Cancer Atlas Program (TCGA) database (https://www.cancer.gov/ccg/research/genome-sequencing/tcga), comprising training set TCGA-HNSC, single-cell RNA sequencing (scRNA-seq) dataset GSE206332, and validation sets GSE59102 and GSE65858. The Table 1 provided detail information of these datasets. Besides, 2118 ac4C modification-related genes (acRGs) were obtained from a previous study[17] (Table S1).

**Table 1.** Information of datasets utilized in this research.

| Dataset | Sample (controls/patients) | Sequencing platform |
| --- | --- | --- |
| TCGA-HNSC | 12/115 | RNA-seq |
| GSE206332 | 3/3 | GPL18573 |
| GSE59102 | 13/29 | GPL6480 |
| GSE65858 | 48 LSCC patients | GPL10558 |

### 2.2 Single-cell RNA sequencing data processing

We first performed quality control of scRNA-seq data to eliminate abnormal cells, including nFeature_RNA greater than 200 and less than 7,000, nCount_RNA larger than 100 and less than 60,000, and percent_MT less than 20. Data normalization was performed using the NormalizeData function in the R package Seurat (v 4.3.0). Subsequently, highly variable genes were identified via the FindVariableFeatures function. Next, we downscaled the data using principal component analysis and clustered them using the FindNeighbors and FindClusters functions. Cellular annotations were performed with the help of published and more classical marker genes.

### 2.3 Cell subpopulation identification and enrichment analysis

Malignant epithelial cells were identified using InferCNV (v1.24.0) by comparing their copy number variation (CNV) levels with those from normal samples. These cells were subsequently clustered to define distinct subpopulations. To explore the functional characteristics of malignant cells, we used the AUCell algorithm to analyze the enrichment of a predefined set of cell stemness-related genes[18]. Differential gene expression across the identified subpopulations was determined using the RunDEtest function, followed by gene ontology analysis to interpret their functional relevance. The threshold for differentially expressed genes was set to |avg_log2FC| > 1 and the adjusted P-value (p_val_adj) < 0.05. Subsequently, we used the AUCell algorithm to evaluate the expression characteristics of acRGs in different cell subpopulations, defining the cell subpopulation with the highest AUC value as the key subpopulation.

### 2.4 Pseudotime analysis, cell communication, and transcription factor prediction analysis

Sequential analysis can be used to infer changes in cells during dynamic processes. We used CytoTRACE and Monocle algorithms to deeply infer and analyze the differentiation and development trajectories of malignant epithelial cells. To further identify key cell subpopulations associated with acRGs, we used the AUCell algorithm to evaluate the expression characteristics of acRGs in cell subpopulations. The communication patterns of key cell subpopulations and related signaling pathways were explored using the CellChat tool (v 1.6.1). The single-cell regulatory network inference and clustering (SCENIC) pipeline was employed to infer transcription factor activity, thereby revealing potential regulatory interactions between different cell subpopulations.

### 2.5 Univariate Cox regression analysis and machine learning

Using the TCGA-HNSC dataset, we selected the top 300 genes with significantly upregulated expression in key cell subpopulations for univariate Cox regression analysis, screening out candidate genes significantly associated with overall survival for further analysis. The candidate genes were subjected to further screening via four distinct machine learning approaches-least absolute shrinkage and selection operator (LASSO) regression, support vector machine-recursive feature elimination (SVM-RFE), extreme gradient boosting (XGBoost), and random forest (RF) algorithms. The LASSO method functions by introducing a penalty term that compresses regression coefficients, effectively filtering for genes with non-zero values. SVM-RFE, XGBoost, and RF algorithms assess the importance of candidate genes and rank them, ultimately retaining the top 20 genes. Key prognostic genes were defined as the intersecting results from the four machine learning algorithms. Differential expression of these genes was further evaluated by comparing tumor and paired normal tissues within the training and external validation datasets. The surv_cutpoint function is used to determine the optimal cut-off point for each prognostic gene, thereby dividing patients into high-expression and low-expression groups. The prognostic value of the identified genes was then assessed using Kaplan-Meier (KM) analysis, evaluating the association between their expression levels and patient survival.

### 2.6 Construction and evaluation of prognostic risk model

Using the machine learning-selected prognostic genes, we developed a risk prediction model via multivariate Cox regression, which was then evaluated by receiver operating characteristic (ROC) curve analysis. Nomogram charts constructed using the R package regplot (v 1.1) were used to predict patient survival rates at 2, 3, and 4 years after diagnosis. Calibration curves were used to validate the accuracy of the model’s predictions. Additionally, we used SHAP analysis to explore the specific contributions of each feature in the risk model to the model’s predictions. To evaluate the prognostic impact of risk stratification, the optimal cutoff for dividing patients was first determined using the fit$baseline output from SHAP analysis, categorizing them into high- and low-risk groups. Subsequently, KM survival analysis was employed to compare survival outcomes between these groups.

### 2.7 Immune infiltration, anti-cancer immune cycle, immune response, and immune subtype analyses

The CIBERSORT algorithm was used to investigate the immune infiltration status of LSCC patients with different risks. To explore the differences in tumor immune cycle scores among patients with different risks, we assessed each patient’s activity score using tumor immune phenotype tracking (TIP, https://biocc.hrbmu.edu.cn/TIP/). In addition, we used the tumor immune dysfunction and exclusion (TIDE) online tool (http://tide.dfci.harvard.edu/) to calculate immune scores for gene expression data from different risk groups to assess immune exclusion and dysfunction in the tumor immune microenvironment. Each subtype reflects a specific pattern of immune response activity in the tumor microenvironment. Therefore, we employed the R package ImmuneSubtypeClassifier (v 0.1.0) to evaluate the distribution of immune subtypes among patients in different risk groups.

### 2.8 Somatic mutations, drug sensitivity, and pan-cancer analyses

LSCC patient somatic mutation profiles were retrieved and characterized using R packages TCGAmutations (v 0.4.0) and maftools (v 2.20.0). Subsequently, drug sensitivity profiles, expressed as half-inhibitory concentration (IC50) values, were predicted from the Drug Sensitivity in Cancer (GDSC) database using the R package pRRophetic (v 0.5). Statistical comparison of IC50 values across risk groups was performed with the Wilcoxon rank-sum test. Additionally, the gene set cancer analysis (GSCA, http://bioinfo.life.hust.edu.cn/GSCA/#/expression) database was used to assess the expression of prognostic genes in other cancers, including bladder urothelial carcinoma (BLCA), stomach adenocarcinoma (STAD), colon adenocarcinoma (COAD), prostate adenocarcinoma (PRAD), esophageal carcinoma (ESCA), thyroid cancer (THCA), head and neck squamous cell carcinoma (HNSC), kidney renal clear cell carcinoma (KIRC), kidney chromophobe (KICH), kidney renal papillary cell carcinoma (KIRP), breast invasive carcinoma (BRCA), liver hepatocellular carcinoma (LIHC), lung adenocarcinoma (LUAD), and lung squamous cell carcinoma (LUSC).

### 2.9 qPCR validation

The prognostic gene expression was clinically validated using five sets of tumour-normal paired samples recruited between 1 June 2025 and 1 December 2025. These samples were initially collected by the First Affiliated Hospital of Nanjing Medical University following ethical approval (No. 2025-SR-579) and written informed consent. RNA extraction was performed using Trizol reagent (Vazyme, China) for tissue lysis, followed by chloroform (Servicebio, China) and isopropanol (Shanghai Trial, China) purification. RNA concentrations were quantified and standardized prior to reverse transcription into cDNA. gDNA Eraser Mix (Albatross Biology, China) is used for reverse transcription to obtain cDNA. Quantitative PCR was carried out with SYBR Master Mix (Albatross Biology, China) and the primers listed in Table 2. The relative expression of target genes was determined via the 2^-ΔΔCT^ method, normalized to the endogenous control β-actin.

**Table 2.**
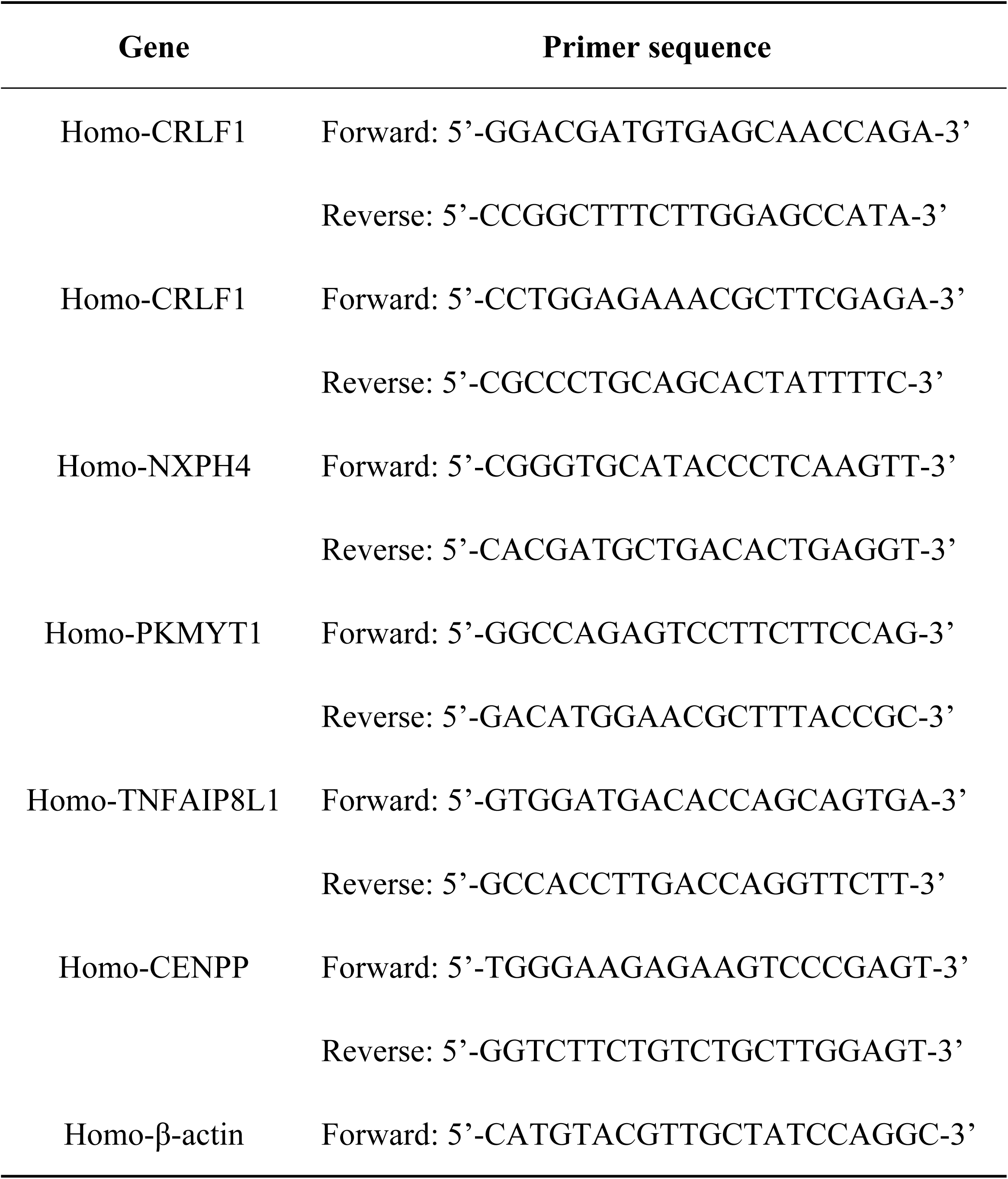
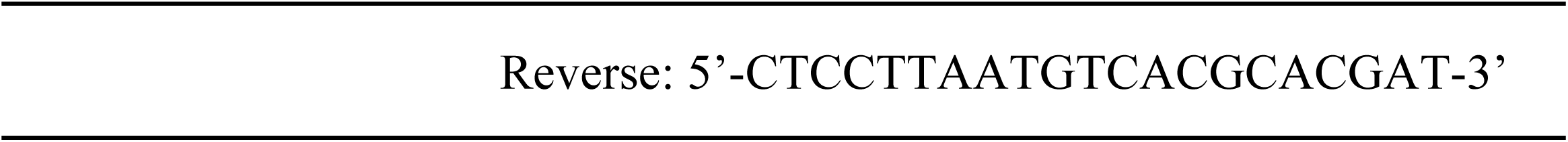
The list of primer sequences.

### 2.10 Statistical analysis

Statistical analysis was performed using R software (v 4.2.2) and GraphPad Prism (v 10.1.2). Network visualizations were generated with Cytoscape software (v 3.9.1). Comparisons between two groups were conducted using Student’s *t*-test for normally distributed data and the Mann-Whitney U test for nonparametric data. A p-value of less than 0.05 was considered statistically significant in all analyses.

## 3. Results

### 3.1 Identification of malignant epithelial cells

We obtained a total of 50,357 cells from the GSE206332 dataset, and after data quality control, 42,937 cells remained (Fig. S1A, Fig. 2A). Based on the top 4,000 highly variable genes selected for further study, 13 clusters were subsequently identified by principal component analysis (Fig. 2B-C, Fig. S1B). Using marker genes, we annotated these 13 clusters, ultimately identifying five cell types, including B cells, epithelial cells (ECs), fibroblasts, myeloid cells, and T cells (Fig. S1C, Fig. 2D-E). Since ECs are the most abundant cell type in both normal and tumor tissues (Fig. 2F), we used ECs from normal samples as a control and performed CNV analysis on ECs from tumor tissues (Fig. 2G-H). The results showed that 14,465 ECs, excluding those in clusters 2 and 4, had high CNV values, which were defined as malignant epithelial cells (MECs) for subsequent analysis.

**Figure 2.**
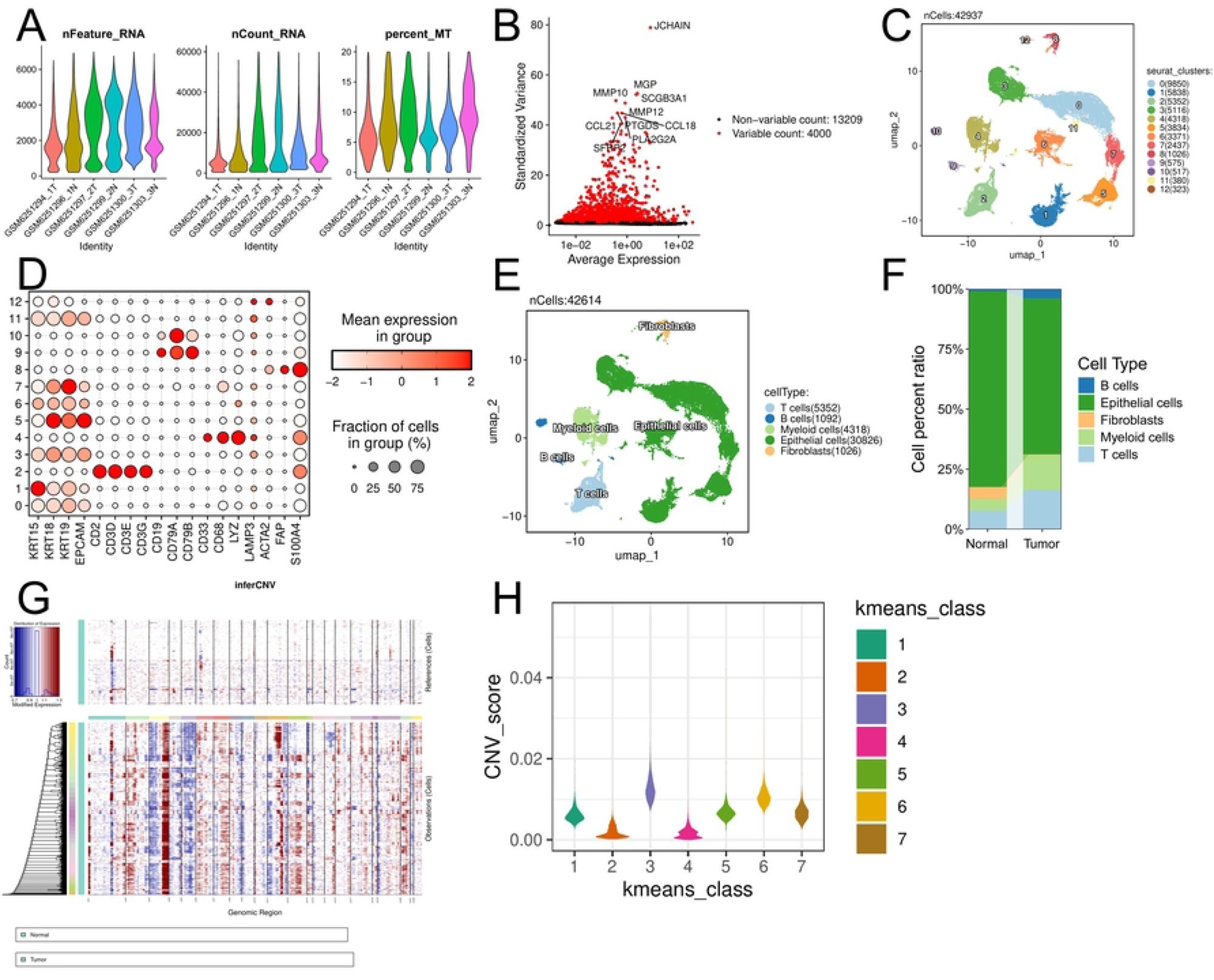
Identification of malignant epithelial cells. (**A**) The nFeature_RNA, nCount_RNA, and percent_MT after quality control. (**B**) Variance plot showing all 17,209 genes, with the top 4,000 highly variable genes highlighted in red. (**C**) UMAP plot showing 13 clusters detected. (**D**) Dot plot showing marker gene expression across the 13 identified cell clusters. (**E**) UMAP plot demonstrating five cell populations identified. (**F**) Cell percentage chart, with epithelial cells accounting for the highest percentage. (**G**) The inferCNV analysis results graph showing the CNV levels of epithelial cells in different samples. (**H**) Violin plot showing the CNV values for each cluster.

### 3.2 Characterization of malignant epithelial cell subpopulations

We conducted an in-depth investigation of MECs, clustering them and identifying five cell subpopulations, named Malig_Epi_0 (MEC0), Malig_Epi_1 (MEC1), Malig_Epi_2 (MEC2), Malig_Epi_3 (MEC3), and Malig_Epi_4 (MEC4) (Fig. 3A). Analysis further revealed that most cells in these subpopulations were arrested in the G1 phase of mitosis (Fig. 3B). By analyzing the biological characteristics of the MECs, we found that MEC3 had the highest stemness AUC value, while MEC2 had the highest G2M score (Fig. 3C, Fig. S2A). To investigate the biological functions of different MEC subpopulations, we first performed differential expression analysis on the five subpopulations to screen for DEGs (Fig. 3D, Fig. S2B). Functional enrichment analysis revealed significant heterogeneity among different MEC subpopulations in key biological processes such as cell cycle regulation, metabolic activity, and epidermal differentiation (Fig. 3E-F, Fig. S2C-E). To further assess the potential developmental trajectories of these subpopulations, we performed pseudo-time analysis. The results indicated that the MEC3 subpopulation had the highest CytoTRACE score, suggesting the strongest stemness (Fig. 3G). By combining CytoTRACE predictions with pseudotime ordering, it was observed that the MEC trajectory evolved directionally from the MEC3 to the MEC1 subpopulation (Fig. 3H-I).

**Figure 3.**
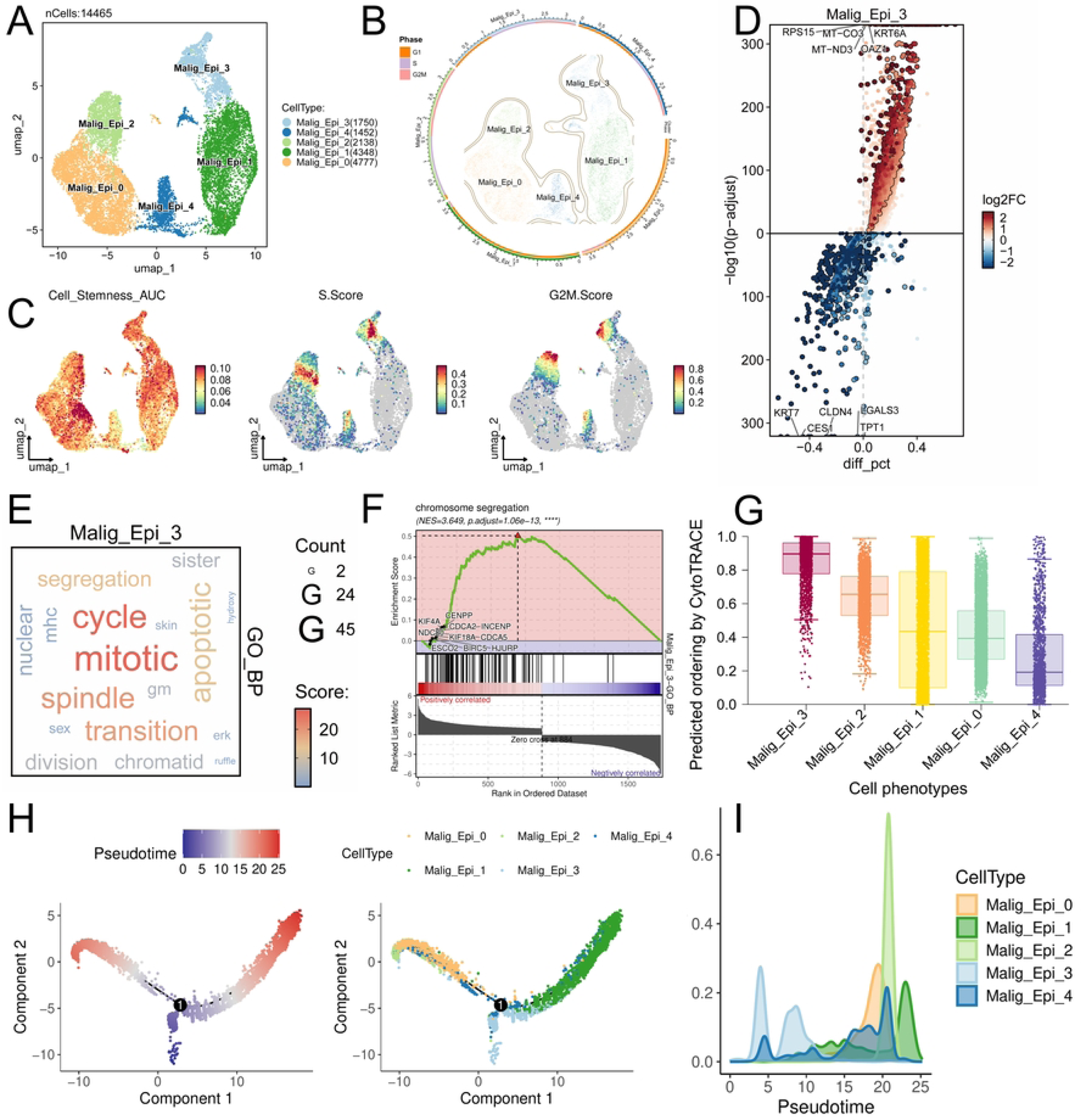
Characterization of malignant epithelial cell subpopulations. (**A**) UMAP plot showing the distribution of the five subpopulations of MECs. (**B**) Circular UMAP plot depicting the distribution of different cell subpopulations across cell cycle stages. (**C**) UMAP plot highlighting key features between subgroups, including Cell Stemness_AUC, S Score, and G2M Score. (**D**) Volcano plot depicting differentially expressed genes in the MEC3 subgroup. (**E**) Word cloud showing the specific biological processes corresponding to the highly enriched genes in the MEC3 subpopulation. Larger font sizes indicate a greater number of enriched genes in that biological process, while darker colors indicate a higher enrichment score for the genes. (**F**) GSEA plot displaying the most significantly enriched pathways in the MEC3 subpopulation. (**G**) The CytoTRACE algorithm evaluating the stemness of each subpopulation. (**H**) Monocle2 differentiation trajectory plot of cell subpopulations. (**I**) Spine diagram displaying the density distribution of cells in each subgroup on the pseudo-time trajectory.

### 3.3 Cell communication between MEC3 and other cell populations

By comparing the AUC scores of each subgroup with the ac4C modification gene set, we found that MEC3 had the highest score and was therefore defined as the key subgroup for subsequent analysis (Fig. 4A). We screened the top five TFs in MEC3 based on AUC scores, including ODC1, FOSL1, IRX2, MYC, and KLF2, for further evaluation. The results showed that the expression levels of these five TFs exhibited significant heterogeneity across different MEC subgroups (Fig. 4B). Additionally, we assessed cell-to-cell communication between MEC3 and other subpopulations. Significant cell-to-cell interactions were observed between MEC3, other subpopulations, and fibroblasts (Fig. 4C). Based on cell-to-cell outgoing and incoming signal results, MIF signaling was significantly enriched in MEC3 (Fig. 4D). A deeper analysis of the MIF signaling pathway revealed that MEC3 is the primary sender of this signal, which is transmitted to myeloid cells (Fig. 4E-F). Specifically, MEC primarily interacts with myeloid cells via MIF-(CD74+CD44) and MIF-(CD74+CXCR4) (Fig. 4G). Collectively, these findings indicate that MEC3 plays an important role in the tumor microenvironment.

**Figure 4.**
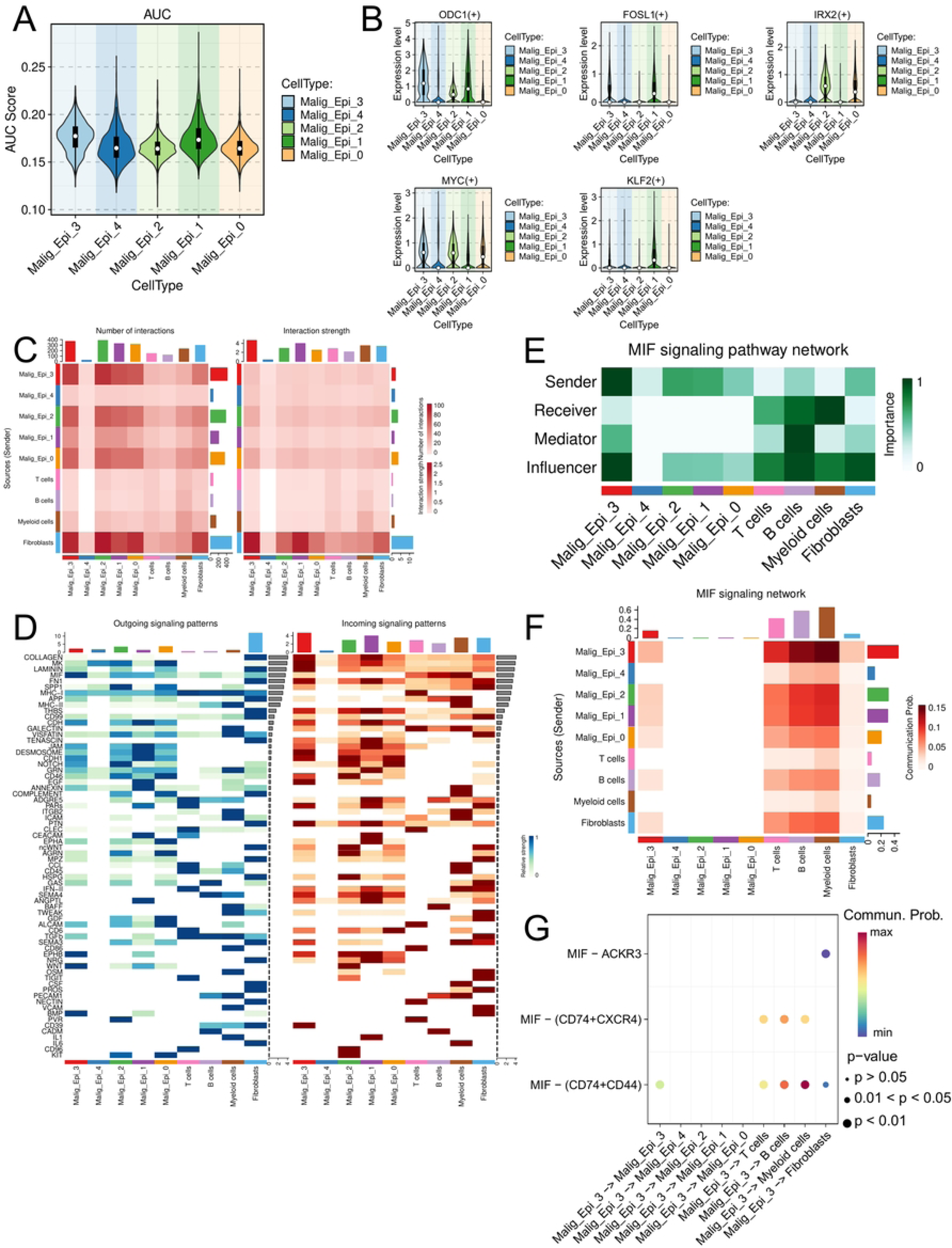
Cell communication between MEC3 and other cell populations. (**A**) AUC scores between five MEC cell subpopulations and the ac4C modification gene set. (**B**) Violin plot showing the expression levels for the top five transcription factors (ranked by AUC score) in each MEC subpopulation. (**C**) Heatmap depicting intercellular interaction frequency (left panel) and interaction strength (right panel), with color intensity proportional to the respective metric. (**D**) Heatmap illustrating the signal transmission patterns between different cell groups, including outgoing signaling patterns (left) and incoming signaling patterns (right). The darker the color, the stronger the signal intensity. (**E**) Heatmap evaluating the strength of the roles played by MIF signals as senders, receivers, regulators, and mediators in various cell types. The darker the color, the greater the involvement. (**F**) Heatmap representing the probability of signal transmission between cells, with darker colors indicating higher probabilities. (**G**) Dot plot revealing the probability of interaction between different ligand-receptor pairs in different cell types. The redder the color of the dots, the greater the probability of communication.

### 3.4 Identification of seven key prognostic genes in LSCC

To identify LSCC-associated prognostic genes, the top 300 most highly expressed genes in the MEC3 subgroup were first selected. Following the selection, univariate Cox regression analysis was performed, which identified 26 genes significantly associated with patient prognosis (Fig. 5A). Next, we used multiple machine learning algorithms to perform a secondary screening of these 26 genes. LASSO regression identified 19 genes with non-zero coefficients (Fig. 5B), while SVM, XGBoost, and RF algorithms each extracted the top 20 genes ranked by feature importance (Fig. 5C-E). A reverse intersection analysis of the candidate genes identified by these four algorithms yielded seven prognostic genes: BARX1, FHL2, NXPH4, PKMYT1, TNFAIP8L1, CRLF1, and CENPP (Fig. 5F).

**Figure 5.**
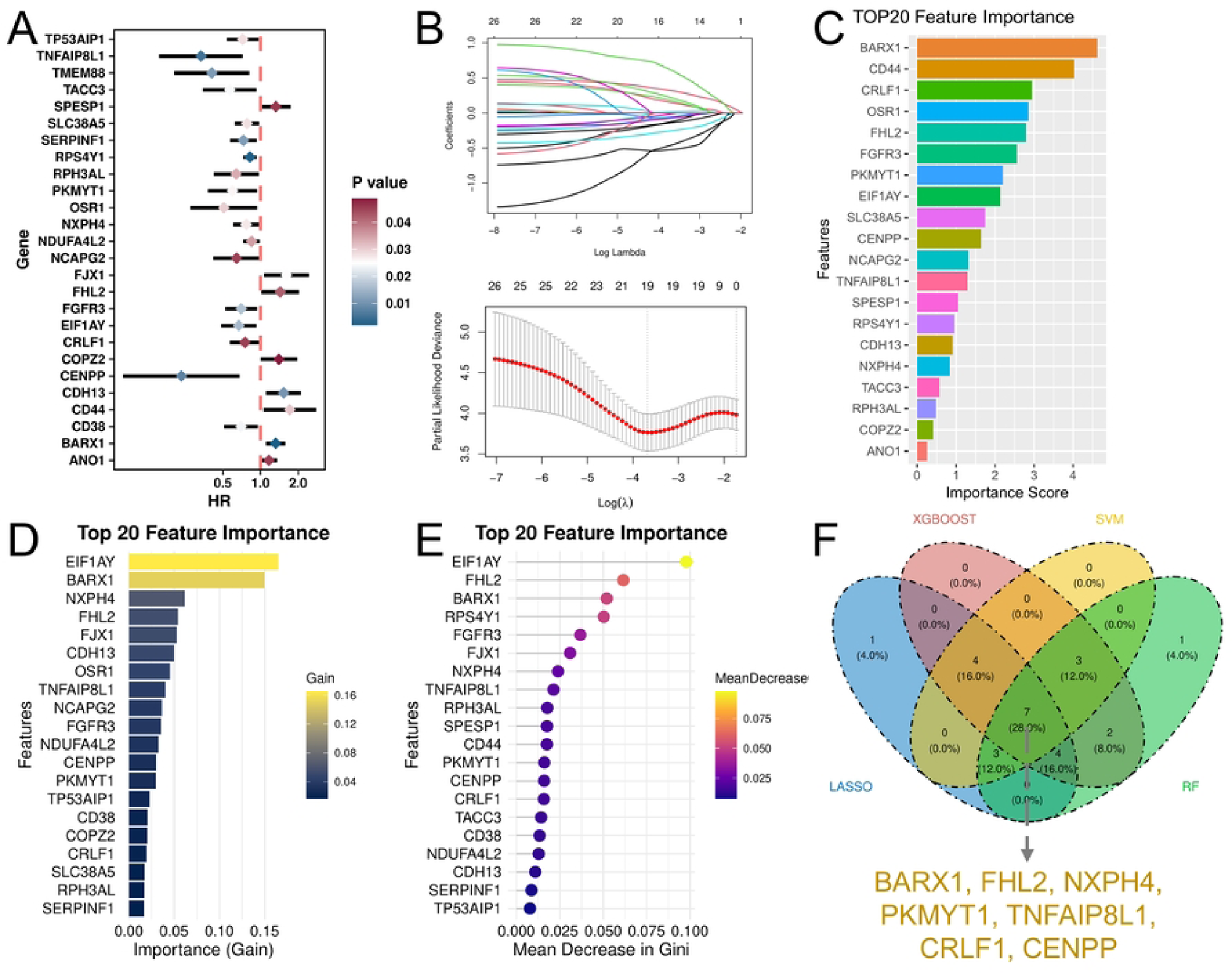
Identification of prognostic genes in LSCC. (**A**) Forest plot showing the results of single-factor Cox regression screening. (**B-C**) Gene selection process for the MEC3 subpopulation gene set, including cross-validation curve (upper) and LASSO coefficient path (bottom). (**C-E**) Top 20 genes ranked by feature importance in SVM (**C**), XGBoost (**D**), and RF (**E**). (**F**) Venn diagram depicting that the four algorithms identified seven genes in common.

### 3.5 Construction and validation of prognostic risk model

Based on the seven prognostic genes identified, we constructed a prognostic risk model. The model formula is: riskScore = 0.45695076 × expression value of FHL2 - 0.04777116 × expression value of CENPP + 0.54713187 × expression value of BARX1 - 0.54279761 × expression value of NXPH4 - 0.27492320 × expression value of PKMYT1 - 0.19990061 × expression value of TNFAIP8L1 - 0.17916632 × expression value of CRLF1. A nomogram was used to visually demonstrate the predictive capability of the model (Fig. 6A). The model was validated using calibration curves, which demonstrated close agreement between predicted and actual values, confirming its high accuracy in predicting survival probability (Fig. 6B). Additionally, ROC results showed that the AUC values for different datasets at different time points were all above 0.7 (Fig. 6C-D), further validating the predictive capability of the model. To further elucidate the contribution of prognostic genes to the model, we used SHAP analysis to rank their importance. The results indicated that BARX1, NXPH4, and FHL2 contributed the most to the model’s predictions (Fig. 6E). It is worth noting that the high expression of BARX1 and FHL2 corresponds to positive SHAP values, suggesting they may be potential risk factors (Fig. 6F). LSCC patients were categorized into high- and low-risk groups based on an optimal cutoff (1.18E-16) identified from SHAP scores. Kaplan-Meier survival analysis demonstrated a significantly more favorable prognosis for the low-risk group (Fig. 6G, Fig. S3A). Additionally, expression analysis showed an upward trend for FHL2 and BARX1 in high-risk patients, contrasting with the enrichment of several other genes in the low-risk group (Fig. 6H, Fig. S3B). The concentration of deaths in the high-risk group further validated the close association between risk scores and patient survival outcomes (Fig. 6I-J, Fig. S3C-D).

**Figure 6.**
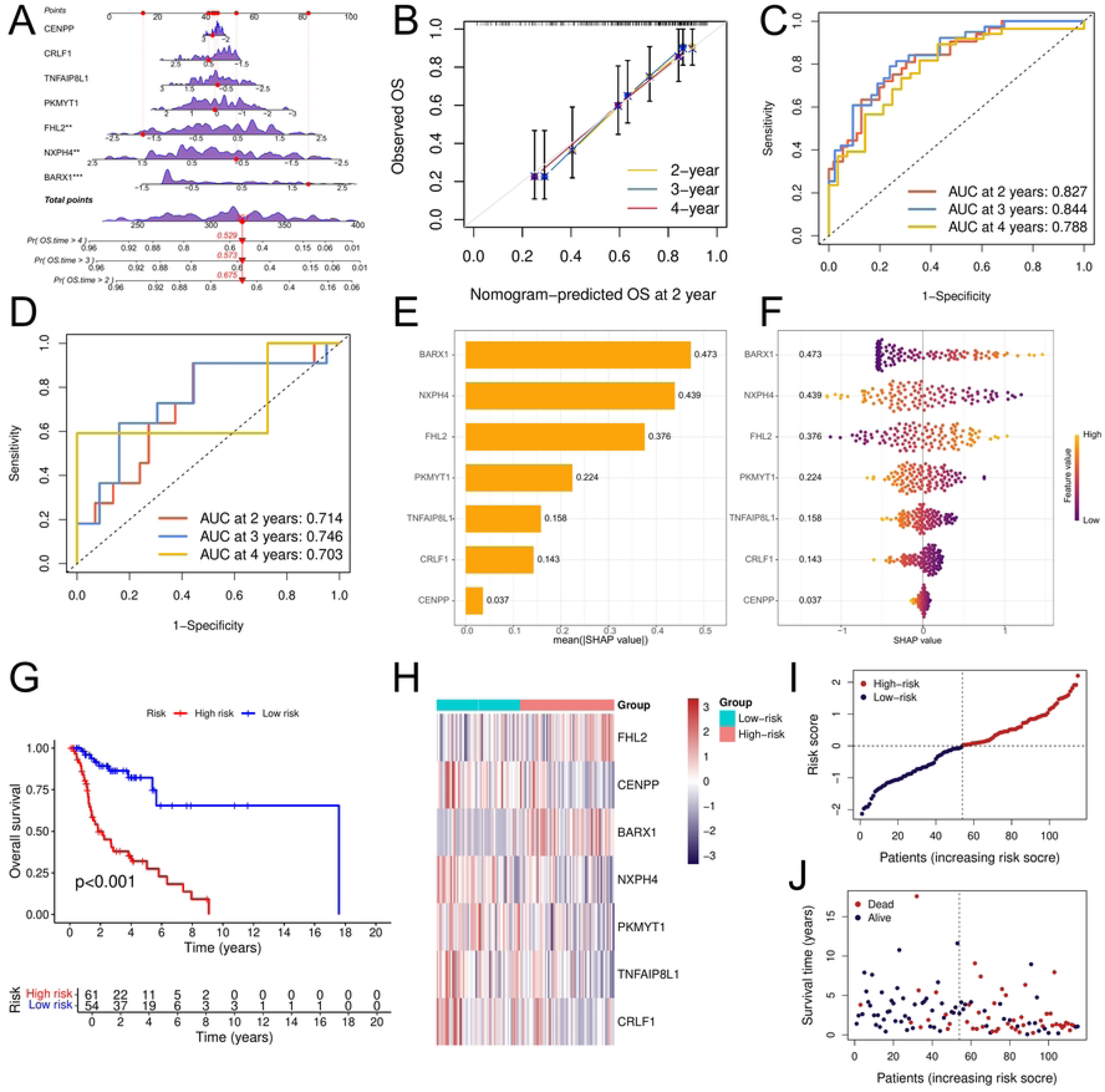
Development of a prognostic model for LSCC. (**A**) Nomogram containing seven prognostic genes is used for risk prediction in LSCC patients. (**B**) Calibration curve showing concordance between predicted and observed outcomes. (**C**-**D**) ROC curves assessing predictive accuracy in the training (**C**) and validation (**D**) sets. (**E**) Bar chart showing the importance ranking of prognostic genes in SHAP analysis. (**F**) Honeycomb plot displaying the relationship between SHAP values and expression levels for prognostic genes. (**G**) Kaplan-Meier curves illustrating the difference in overall survival between high-risk and low-risk patient groups. (**H**) Heatmap of prognostic gene expression trends across different risk groups. (**I**) Comparison of the prognostic risk scores between the two patient groups. (**J**) Distribution of patient survival status, stratified by risk group (separated by dashed lines).

### 3.6 Association between prognostic gene expression and survival in LSCC patients

We further evaluated the expression levels of prognostic genes in LSCC patients using the dataset. The analysis revealed a significant downregulation of CRLF1 in tumor tissues (p < 0.05), while FHL2 exhibited no significant differences in either the training or validation sets (Fig. 7A-B). All five remaining genes were significantly upregulated in tumors (p < 0.05). The findings were validated independently via qPCR on clinical samples, testing all prognostic genes except FHL2, which had shown no significant differences previously (Fig. 7C). The results confirmed highly significant differential expression for all tested genes between the two groups (p < 0.0001). However, with the exception of BARX1, all others aligned with the expression trends observed in the dataset, further validating the reliability of the bioinformatics analysis findings. The association between gene expression levels and patient survival was further evaluated. Survival analysis revealed that patients with lower expression of BARX1 and FHL2 had significantly better survival rates compared to those with higher expression (p < 0.0001, Fig. 7D). Conversely, elevated expression of the remaining five genes was associated with more favorable survival outcomes (p < 0.01). In summary, these findings further support the notion that prognostic genes may be involved in the pathological mechanisms of LSCC.

**Figure 7.**
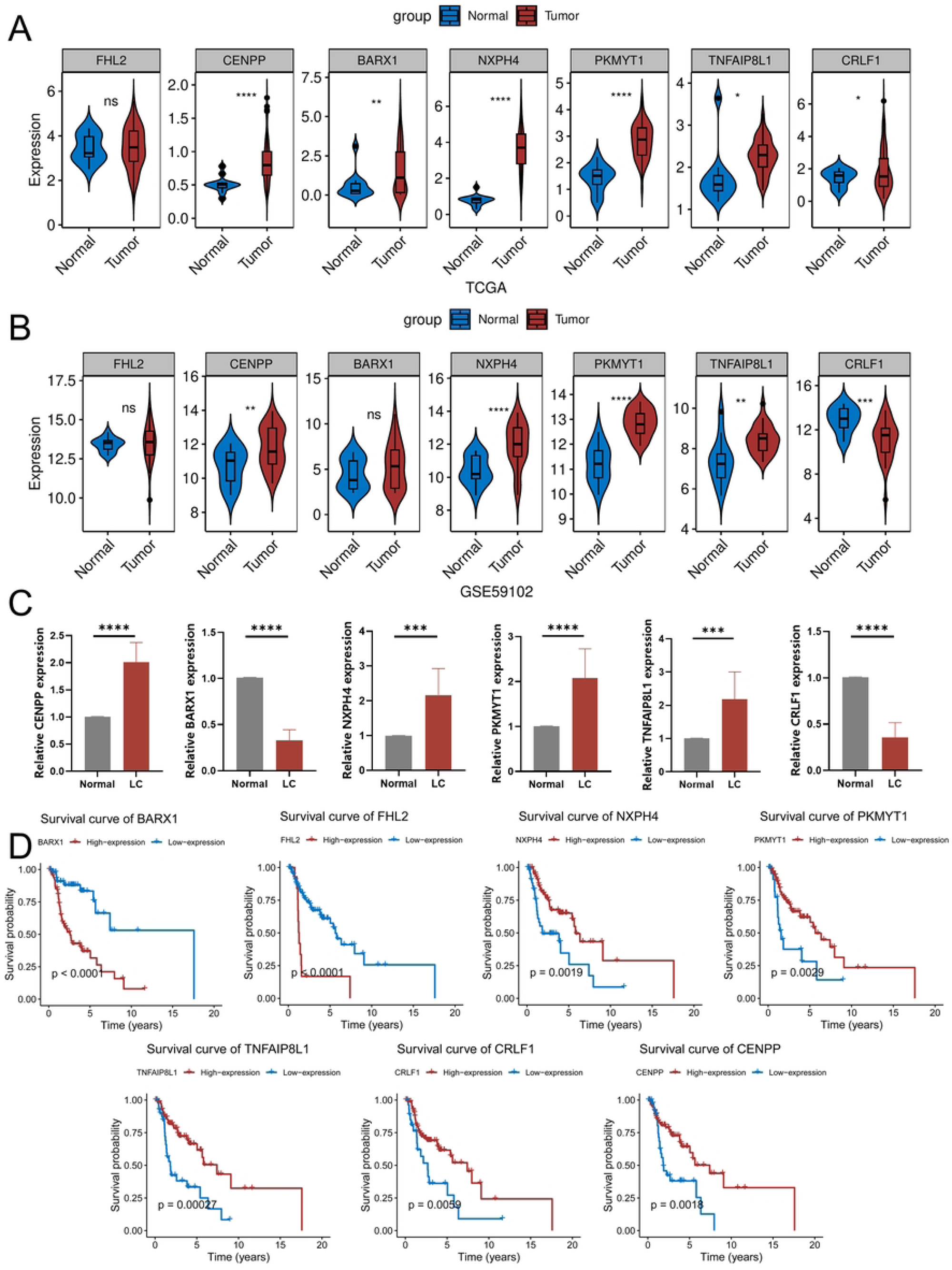
Prognostic gene expression and survival analysis. (**A-B**) Violin plots comparing prognostic gene expression between tumor and normal tissues in the TCGA-HNSC (**A**) and GSE59102 (**B**) datasets. (**C**) Bar chart showing the expression patterns of prognostic genes in clinical samples. (**D**) Kaplan-Meier survival curves showing the association between prognostic gene expression levels and patient survival. ns > 0.05, *p < 0.05, **p < 0.01, ***p < 0.001, ****p < 0.0001.

### 3.7 Immune cell infiltration imbalance in LSCC patients

To explore potential responses to immune checkpoint inhibitors, TMB was analyzed in all training set samples. Most samples in both risk groups harbored gene mutations, with TP53 and TTN being particularly frequent (Fig. 8A). Next, we assessed immune cell infiltration in patients with different risk levels. Profiling of immune cell infiltration indicated that M0 and M2 macrophages were the most abundant subsets (Fig. 8B). A comparative analysis further identified activated memory CD4+ T cells as differentially enriched in the low-risk group and M0 macrophages in the high-risk group (p < 0.01, Fig. 8C), implying a connection to immune evasion. Additionally, correlation analysis results indicated that BARX1, CENPP, FHL2, NXPH4, and PKMYT1 were significantly correlated with immune cell infiltration levels (Fig. 8D), suggesting that these genes may influence LSCC by regulating immune responses.

**Figure 8.**
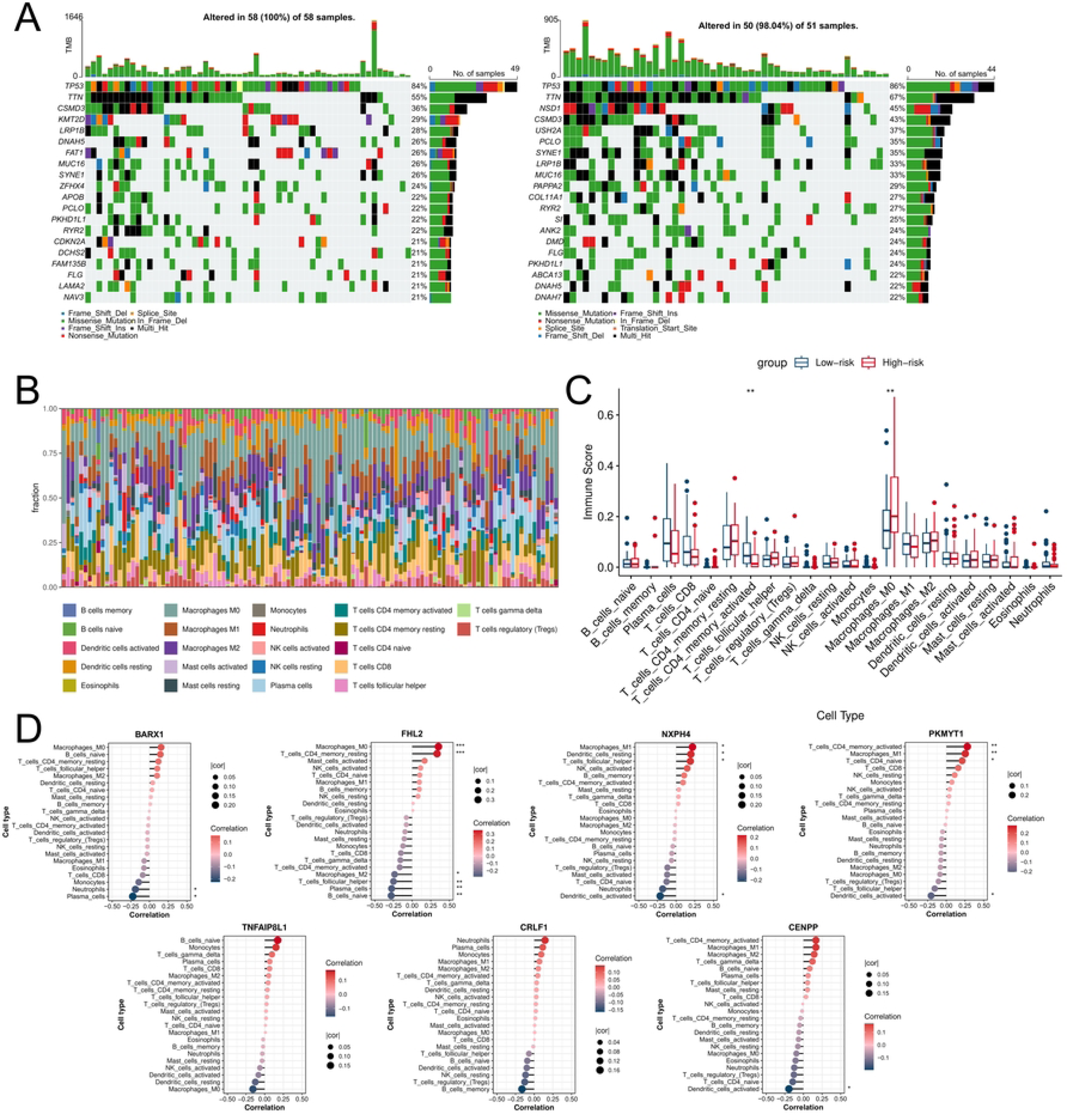
Imbalance of immune cell infiltration in LSCC. (**A**) Somatic mutation profiles for patients in the high-risk (left) and low-risk (right) groups, showing gene mutations across samples and color-coded by mutation type. (**B**) Immune cell abundance bar chart showing the infiltration abundance of 22 types of immune cells in each sample. (**C**) Box plot comparing immune scores of 22 immune cell types between patient risk groups. (**D**) Lollipop plot showing the correlation between seven prognostic genes and 22 types of immune cells. *p < 0.05, **p < 0.01, ***p < 0.001.

### 3.8 Alterations in the immune landscape of LSCC patients

To evaluate the anti-tumor immune status across risk groups, we analyzed the anti-cancer immunity cycle analysis. Step 5 (immune cell tumor infiltration) was the only phase significantly enriched in high-risk patients (p < 0.05, Fig. 9A), indicating a more active immune response and greater infiltration capacity in this group. Immunotherapy response analysis showed that the high-risk group had higher Exclusion scores, while the low-risk group had higher Dysfunction scores (p < 0.05, Fig. 9B). This finding suggests that tumors in high-risk patients are more likely to evade immune surveillance by excluding immune cells. Additionally, we classified tumor samples in the training set and found only two subtypes: C1 (wound healing) and C2 (IFN-γ dominant) (Fig. 9C). Furthermore, drug sensitivity analysis revealed pronounced sensitivity differences between risk groups. Specifically, the high-risk group demonstrated significantly lower IC50 values for Bleomycin, BMS-708163, Gemcitabine, NSC.87877, and Roscovitine compared to the low-risk group (p < 0.0001; Fig. 9D), indicating enhanced efficacy for these patients. Conversely, the remaining five agents showed significantly lower IC50 values in the low-risk group (p < 0.0001), suggesting greater suitability for this patient subset. To assess the expression of prognostic genes across different cancers, we conducted a pan-cancer analysis (Fig. 9E). These prognostic genes showed significant upregulation in multiple cancers, indicating a possible important role in tumor biology beyond LSCC.

**Figure 9.**
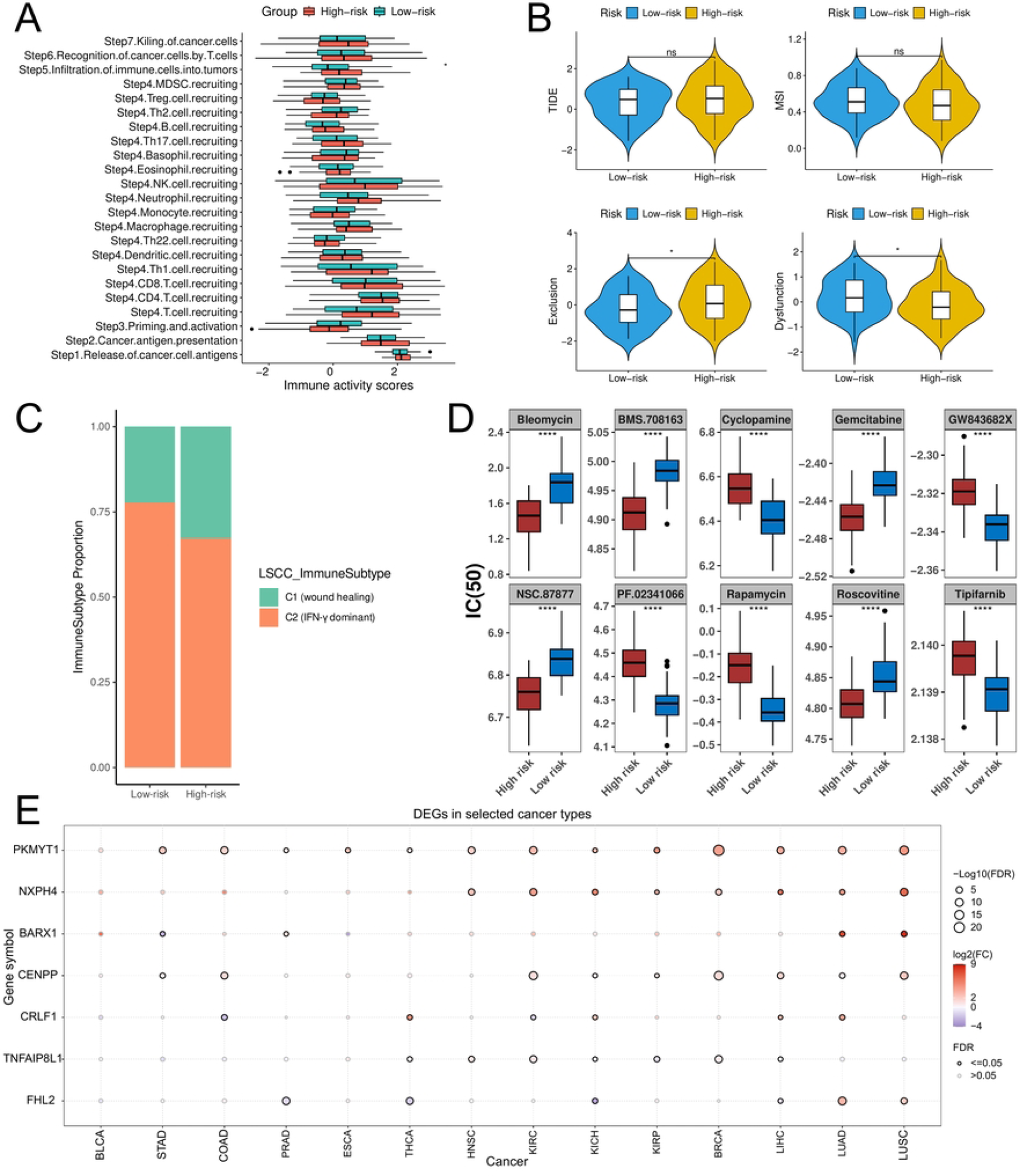
Alterations in the immune landscape of LSCC. (**A**) Box plot depicting immune activity scores at different stages of the anti-cancer immune cycle for high- and low-risk patient groups. (**B**) Violin plot visualizing key immunotherapy response scores (TIDE total, MSI, Dysfunction, Exclusion) for each risk group. (**C**) Bar chart illustrating the proportion of immune subtypes among patients in different risk groups. (**D**) Box plot comparing IC50 values for ten anticancer drugs between risk groups. (**E**) Bubble chart showing the expression of prognostic genes in 14 types of cancer. *p < 0.05, ****p < 0.0001.

## 4. Discussion

The prognosis of LSCC remains poor, with a 5-year survival rate markedly lower than that of many other malignancies[1]. Because ac4C modification drives tumor initiation and progression in multiple cancer types, clarification of its role in LSCC may open unexplored therapeutic avenues[19]. Here, by mining scRNA-seq data we discriminated malignant epithelial cells from non-malignant compartments and resolved them into five transcriptionally distinct sub-clusters. Among these, the MEC3 subpopulation displayed the strongest enrichment for acRGs and was therefore chosen for downstream analyses. Integrating MEC3-specific signatures with bulk transcriptomes identified seven robust prognostic genes-BARX1, FHL2, NXPH4, PKMYT1, TNFAIP8L1, CRLF1 and CENPP-whose collective behavior strongly shaped the tumor immune microenvironment (TIME) of LSCC.

Using the scRNA-seq dataset GSE206332, we identified five major cell lineages and verified 14,465 malignant cells through CNV scoring. AUCell analysis further highlighted the epithelial subpopulation MEC3 was strongly associated with ac4C modification. Cell-cell communication analysis revealed that MEC3 serves as the primary source of MIF signaling, which is predominantly received by myeloid cells. We also identified ODC1 and MYC as potential transcriptional regulators of MEC3, suggesting their involvement in modulating LSCC progression. ECs form the structural and functional core of the mucosal barrier and glandular system, supporting essential barrier, secretory, and absorptive functions[20]. Prior studies demonstrate that epithelial-mesenchymal transition can facilitate LSCC invasion and metastasis[21, 22].

MIF functions through the activation of MAPK and PI3K signaling to regulate cellular processes including proliferation, differentiation, and survival, thereby promoting the development of various cancers such as head and neck squamous cell carcinoma and lung carcinoma[23]. Loss of MIF in breast cancer models enhances pro-inflammatory macrophages and diminishes regulatory T cells and tumor-associated neutrophils[24]. Meanwhile, ODC1 occupies a central role in polyamine metabolism under c-Myc regulation, impacting biosynthesis and gene expression[25], and its deficiency suppresses tumor progression in non-small cell lung cancer by promoting autophagy and inhibiting proliferation[26]. MYC, an established oncogenic driver, regulates transcriptional programs central to tumor initiation and progression[27]. In preclinical models of head and neck squamous cell carcinoma, MYC inhibition has been demonstrated to curtail cellular proliferation and migration[28]. Together, these findings suggest that MEC3-associated epithelial regulatory networks may play an important role in shaping LSCC pathogenesis and immune interactions.

Subsequently, we identified seven prognostic genes associated with ac4C in MEC3 through a combination of univariate Cox regression analysis and multiple machine learning algorithms. BARX1, a homeobox transcription factor governing embryogenesis and differentiation, promotes EMT in oral squamous-cell carcinoma via Wnt/β-catenin activation and correlates with nodal metastasis[29]. Conversely, a 2022 study demonstrated that knocking out BARX1 increases the sensitivity of lung adenocarcinoma cells to cupid’s arrow syndrome[30], suggesting that the role of BARX1 in LSCC warrants further investigation. As a scaffold protein characterized by four-and-a-half LIM domains, FHL2 plays a key role in bridging signal transduction and transcriptional regulation[31]. In head-and-neck squamous-cell carcinoma it amplifies TGF-β signaling to foster metastasis and may additionally modulate YAP/TAZ-driven proliferation and chemoresistance[32]. NXPH4, a secreted neurexophilin glycoprotein, has not been reported in LSCC but facilitates perineural invasion in pancreatic cancer through the semaphorin-plexin axis[33], suggesting a similar pro-invasive role. PKMYT1, a WEE-family kinase that negatively regulates CDK1 and enforces the G2/M checkpoint, confers genomic instability and cisplatin resistance in esophageal squamous-cell carcinoma[34], analogous DNA-repair defects may underlie therapeutic refractoriness in LSCC. TNFAIP8L1 is characterized as an anti-apoptotic member of the TNFAIP8 family, which modulates caspase activity and contributes to immune homeostasis[35]. In lung cancer, TNFAIP8L1 promotes T-cell exhaustion through PD-L1 up-regulation and positively correlates with M2-macrophage infiltration, implying immune-evasive properties in LSCC[36]. CRLF1, defined as a cytokine receptor-like factor and co-receptor for thymic stromal lymphopoietin, recruits regulatory T cells via JAK2-STAT3 signaling in melanoma and may sculpt an immunosuppressive TIME in LSCC[37]. CENPP, a centromere protein ensuring accurate chromosome segregation, is deleted in hepatocellular carcinoma, leading to aneuploidy[38]. The reduced CENPP expression in LSCC may likewise fuel genomic heterogeneity and progression. Collectively, these genes may converge on pathways that influence tumor progression and immune escape in LSCC.

Immune infiltration profiling revealed profound TIME remodeling. The seven-gene signature correlated tightly with multiple immune subsets, most prominently memory CD4+ T cells and M0 macrophages. Memory CD4+ T cells, central to adaptive immunity, endow the host with long-lived antigen recall and co-ordinate secondary responses. In head-and-neck cancer their abundance predicts favorable outcome[29], presumably by activating cytotoxic T lymphocytes and promoting humoral immunity. However, the LSCC milieu can drive their exhaustion or skew them toward Th2/Treg phenotypes, thereby attenuating anti-tumor immunity[32]. As unpolarized precursors with plastic potential, M0 macrophages are associated with a poor prognosis in oral squamous-cell carcinoma[33]. Within the LSCC niche they can be polarized to the pro-tumorigenic M2 phenotype, which secretes TGF-β and other cytokines that suppress T-cell function while fostering angiogenesis and metastasis [34]. In risk stratification for laryngeal cancer, the high-risk group exhibited higher immune exclusion scores, suggesting their tumor microenvironment may achieve immune escape by preventing immune cell infiltration[39]. Conversely, the low-risk group demonstrated higher immune dysfunction scores, indicating that while immune cells can infiltrate tumor tissue, their effector functions remain suppressed-potentially linked to T-cell exhaustion states[40]. These findings suggest that heterogeneous immune escape mechanisms may underlie differential therapeutic responses in LSCC.

Drug sensitivity analysis indicated that patients stratified by risk level may benefit from distinct therapeutic strategies. Specifically, Gemcitabine and Roscovitine exhibited higher sensitivity in the high-risk group, whereas Cyclopamine and Tipifarnib showed greater efficacy in the low-risk group. Gemcitabine, a pyrimidine antimetabolite widely used in clinical oncology, has demonstrated robust antitumor activity with manageable side effects[41]. Roscovitine, a Cdk5 inhibitor, has been reported to suppress tumor immune evasion in brain-metastatic tumor models[42]. In contrast, Cyclopamine, an inhibitor of Hedgehog signaling, enhances apoptosis and suppresses proliferation when combined with circularly permuted TRAIL in human myeloma cells[43]. Tipifarnib, a farnesyltransferase inhibitor currently under clinical evaluation for multiple malignancies, has also been shown to modulate platelet activity[44]. Collectively, these findings suggest that risk-adapted drug selection may offer a rational and potentially effective therapeutic approach for LSCC.

This study delineates the contribution of ac4C modification to LSCC pathobiology and proposes both prognostic and therapeutic biomarkers. Limitations should, however, be acknowledged. Analyses are confined to publicly available datasets that may carry inherent population bias, and modest sample sizes could inflate false-discovery risk. Second, although qPCR validation supported key bioinformatic findings, ac4C modification itself was not directly quantified, and comprehensive *in vitro* and *in vivo* functional studies will be required to further clarify the mechanistic roles of the identified genes and to assess their translational potential.

## 5. Conclusion

In summary, this study integrates single-cell and bulk transcriptomic analyses to explore the involvement of ac4C-related genes in laryngeal squamous cell carcinoma. Seven prognostic genes-BARX1, FHL2, NXPH4, PKMYT1, TNFAIP8L1, CRLF1, and CENPP-were identified and demonstrated stable prognostic value. In addition, significant remodeling of the tumor immune microenvironment was observed, with high-risk patients exhibiting features of immune exclusion. Together, these findings provide new insights into ac4C-associated regulatory patterns in LSCC and may support improved risk stratification.

## Supplement materials

**Fig S1.**
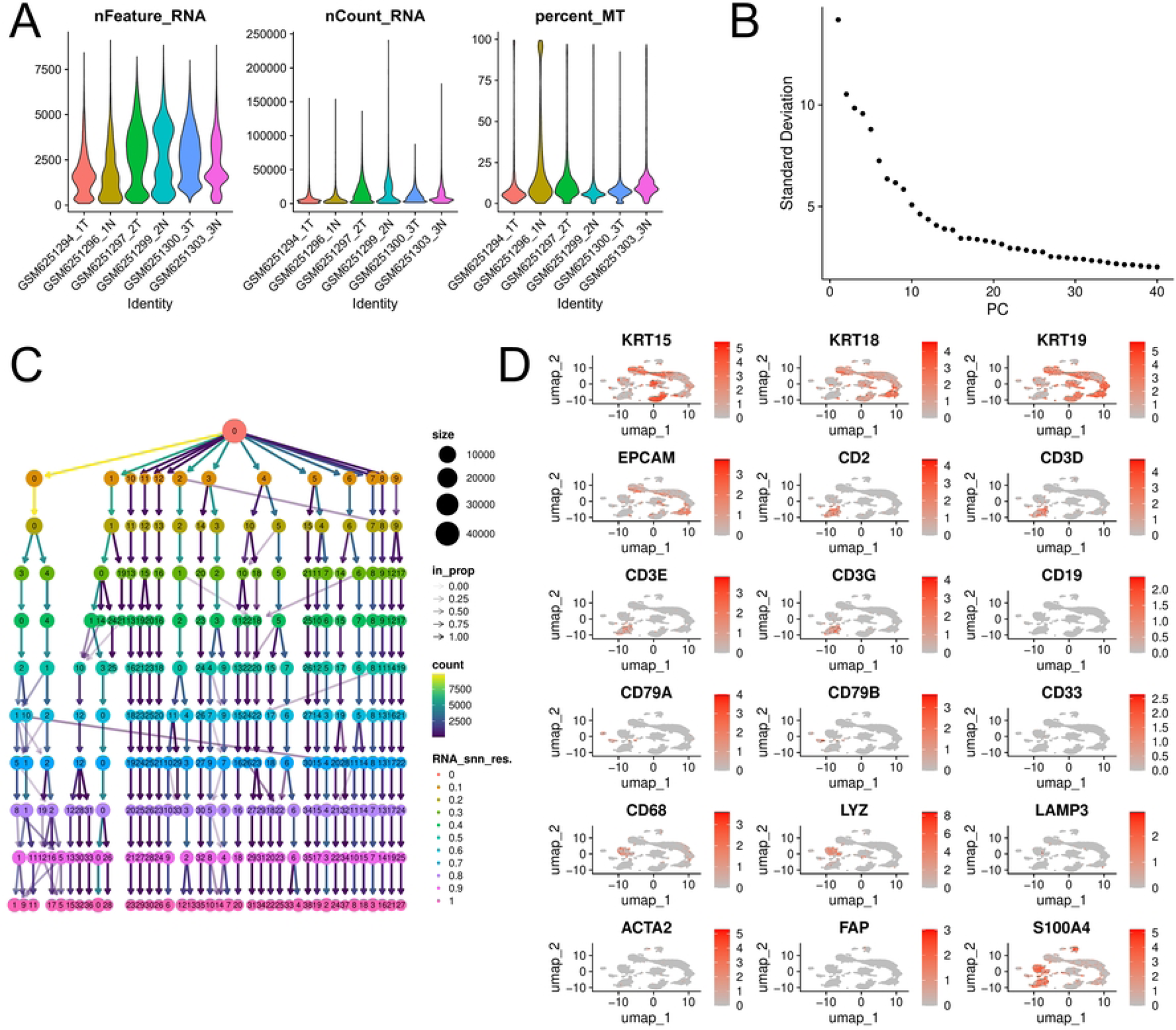
(**A**) The nFeature_RNA, nCount_RNA, and percent_MT before quality control. (**B**) Elbow plot for determining the optimal number of principal components (PCs=30) for clustering. (**C**) Cluster dendrogram displaying the 13 identified cell clusters. (**D**) UMAP plot showing the expression of the marker gene in 13 cell clusters.

**Fig S2.**
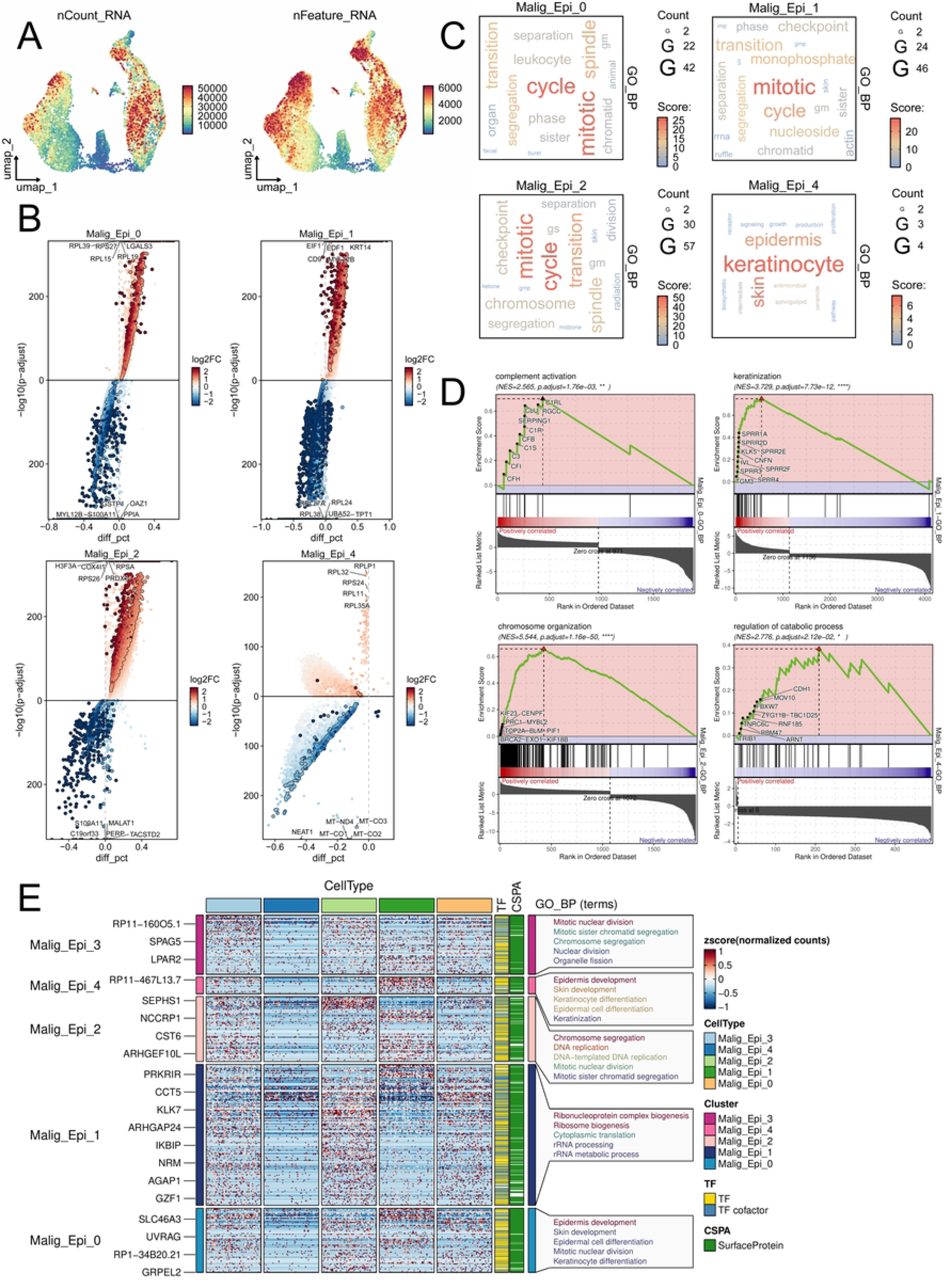
(**A**) UMAP visualization of key technical metrics (nCount_RNA and nFeature_RNA) across cell subpopulations. (**B**) Volcano plot identifying differential genes in the MEC0, MEC1, MEC2, and MEC4 subpopulations. (**C**) Word cloud showing the specific biological processes corresponding to the highly enriched genes in the MEC0, MEC1, MEC2, and MEC4 subpopulations. Larger font sizes indicate a greater number of enriched genes in that biological process, while darker colors indicate a higher enrichment score for the genes. (**D**) GSEA plots displaying the most significantly enriched pathways in the MEC0, MEC1, MEC2, and MEC4 subpopulations. (**E**) Heatmap integrating gene expression patterns of each MEC subpopulation with their top five associated GO BP.

**Fig S3.**
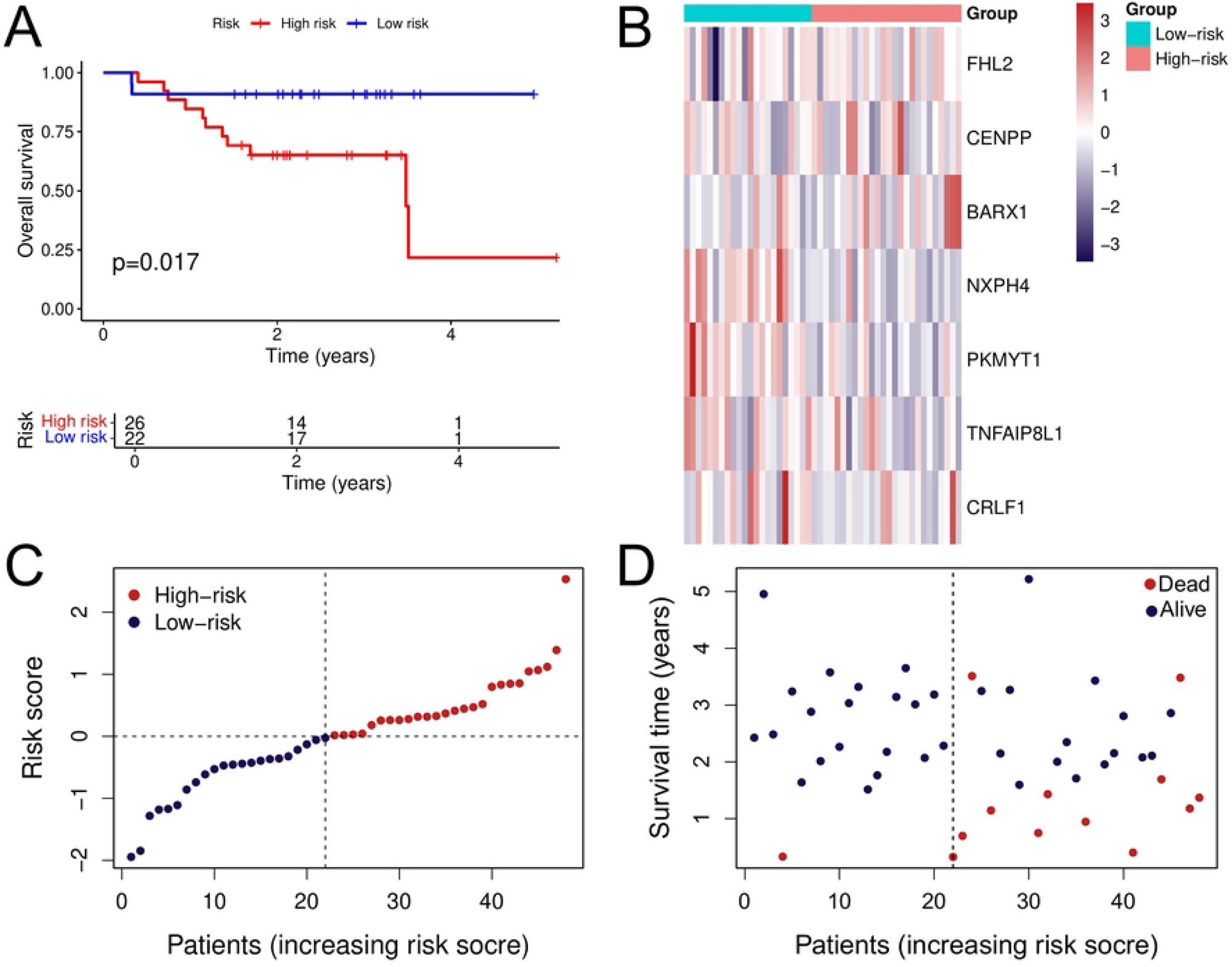
(**A**) KM survival curves showing the survival differences between different risk groups in the validation set samples. (**B**) Heatmaps depicting prognostic gene expression profiles across risk groups within the validation cohort. (**C**) Comparison of model-derived risk scores between high- and low-risk patients in the validation set. (**D**) Comparison of survival status among patients with different risk levels in the validation set.

**Table S1**. The list of ac4C modification -related genes (acRGs).

**Table S2**. The list of genes obtained from LASSO regression, SVM, XGBoost, and RF.

## Author contributions

Conceptualization, L.Q.W. and Y.W.S.; methodology, X.Y.G., J.H.L., and Z.D.L.; software, D.H.C. and J.H.L.; validation, D.H.C., X.C.N., and R.J.Y.; investigation, X.C.N., H.Z., and L.Q.W.; resources, H.Z. and Y.W.S.; data curation, X.Y.G., and R.J.Y.; visualization, J.H.L. and Z.D.L.; supervision, Y.W.S.; writing-original draft preparation, L.Q.W. and X.Y.G.; writing-review and editing, Y.W.S. All authors have read and agreed to the published version of the manuscript. L.Q.W. X.Y.G. D.H.C. X.C.N. H.Z. J.H.L. R.J.Y. Z.D.L. Y.W.S. Liqin Wanga*, Xiaoyang Gonga*, Donghui Chena, Xi chena, Han Zhou, Jiuhuang Lana, Renjing Yea, Zhuoding Luoa, Yawen Shia#

## Institutional review board statement

The study was conducted in accordance with the Declaration of Helsinki, and approved by Research Ethics Committee of The First

Affiliated Hospital with Nanjing Medical University (Approval No. 2025-SR-579).

## Informed consent statement

Informed consent was obtained from all subjects involved in the study.

## Data availability statement

All the RNA-sequencing data and single-cell sequencing data of laryngeal squamous cell carcinoma patients were acquired from The Cancer Genome Atlas (TCGA) (https://www.cancer.gov/ccg/research/genome-sequencing/tcga) and Gene Expression Omnibus (GEO) (http://www.ncbi.nlm.nih.gov/geo/).

## Acknowledgments

Not applicable.

